# Sound Advice: A calibration framework for defining detection space in Passive Acoustic Monitoring

**DOI:** 10.64898/2026.05.20.726556

**Authors:** Prabhat Sharma, Kunapareddy Kezia, Kadaba Shamanna Seshadri

## Abstract

Passive Acoustic Monitoring (PAM) has emerged as a transformative tool for biodiversity assessment in recent years. Despite widespread acceptance and application for conservation-related outcomes, the synergistic effects of hardware limitations, signal propagation, and environmental conditions on how far a signal can be reliably detected remain critically understudied. We quantified changes in signal detectability using Autonomous Recording Units (ARUs) in a tropical agroecosystem using playback experiments of standardised pure-tone (1–8 kHz) in fallow rice paddy fields. We deployed a four-ARU array and broadcast signals over a 50– 300 m distance gradient, and modelled operative detectability of signals using a binomial Generalised Linear Mixed-effects Model (GLMM). Our findings show that the ‘detection space’ of an ARU is highly frequency-dependent and environmentally modulated. Detection probability for low-frequency signals (1 kHz) decreased rapidly (50% threshold at ∼100 m), whereas mid-range frequencies (4–6 kHz) occupied an acoustic window that remained reliably detectable up to 250 m. Higher relative humidity significantly enhanced overall detection, while increasing temperatures disproportionately reduced low-frequency detectability. The orientation of the ARU to the signal source was important as the detection probability declined from 81% for recorders facing the source (0°) to 14% for rear-facing units (180°). Our findings underscore the importance of determining the detection space before undertaking PAM. We propose a ‘Decision Support Framework’ that provides a pathway for researchers to integrate focal taxa traits with technical constraints to determine detection space and optimise study designs when using PAM for monitoring biodiversity and assessing conservation action.

## 1 Introduction

### 1.1 Biodiversity loss and the need for monitoring

Biodiversity is declining globally, resulting in cascading effects on ecosystem function and human wellbeing (1,2). Several countries have ratified biodiversity conservation targets and have pledged to fulfil the Sustainable Development Goals (3,4). Monitoring biodiversity has thus become an important criterion for evaluating progress towards achieving these targets across ecosystems, especially those that have been restored (5). However, long-term datasets of biodiversity remain sparse, especially in the global south, where biodiversity and human populations are high (6,7). Long-term monitoring requires substantial resources, such as sustained funding, field effort, and scientific expertise for physically surveying landscapes for focal taxa over several years, often among the major impediments in the Global South (8).

Technological advancements over the last two decades have led to widespread automation of the monitoring of biodiversity. Advances in the form of improved quality satellite data, longer-lasting batteries, and miniaturisation of devices have increased dramatically, and conservation practitioners have been quick to take advantage (9). Among these advancements is the development of robust Autonomous Recording Units (ARUs). The ARU records acoustic signals at programmed intervals, and the presence or absence of wildlife in each area can be determined. Although researchers have used tape recorders in the past, the ARU is a robust system with improved power consumption, storage, programmability, and analytical workflows, which allow for long-term deployment and scaling of monitoring efforts globally. The modern ARU devices are rugged, durable, and provide enhanced flexibility in programming and storage of data, enabling passive acoustic monitoring (PAM) of biodiversity. Unlike physical surveys conducted by observers, PAM produces reproducible datasets that can be revisited, shared, and analyzed as new algorithms are available (10). Thus, PAM has emerged as a non-invasive and cost-effective technique for monitoring biodiversity across a variety of ecosystems ranging from the open ocean to the forest canopy (11,12). Recent experimental evidence supports this widespread use by demonstrating that PAM can match or outperform human surveys for detecting vocalizing species under varied conditions, depending on observer training and survey design (13,14).

PAM has emerged as a scalable option and is increasingly used to address a wide range of questions in ecology and conservation (11). The widespread use has also necessitated standardized data collection workflows and analytical frameworks (10,15,16). The need for standardization arises from the fact that acoustic signals degrade over distance, across recording devices, and in the environment (17,18). While most ARUs are omnidirectional, very few studies determine the detection space beyond which a signal cannot be reliably detected. This ‘detection space’ or ‘active space’ varies with signal frequency, amplitude, habitat structure, and weather (19). For instance, researchers failed to reliably detect the vocalizations of the koala (*Phascolarctos cinereus*) beyond 100 m in forested habitats but not in the open habitats (20). Therefore, deploying PAM without accounting for this space will likely impact the quality of data and hamper the inter- and intra-site comparisons.

Various methods have been used to determine the detection space of recorders at a site and range from playing back synthesised acoustic signals (19), correlating signals with telemetry data (21), and analysing vocalisations of captive birds, placed at different distances to the recorder (22). Recently, Entling et al. (23) demonstrated the influence of the habitat matrix and the angle of incidence of signals on the detection space of PAM in forested and open agricultural areas. Thus, the efficient deployment of PAM rests on prior knowledge about the study system and must be informed by the detection space within the focal habitats.

### 1.2 Monitoring megadiversity hotspots

Over the last decade, the use of PAM has been rapidly expanding in megadiverse regions such as the Western Ghats in India. The widespread application of PAM ranges from monitoring amphibians in remote and inaccessible habitats such as the forest canopy (24); quantifying avian community structure in the dry and wet grasslands (25); biodiversity monitoring over extended periods across different locations in dry forests (26); determining the presence of rare and endangered birds (27) and recently, PAM was used for evaluating the effectiveness of ecological restoration (28,29). While all these studies provided novel insights, only some of them determined the detection or active space of the recorders, and failing to do so will affect the inferences drawn from such studies, because detected taxa may be present in the larger landscape rather than in the habitat or site being surveyed. For instance, Seshadri and Ganesh (24) deployed paired ARUs in the canopy and on the ground and found that the ARUs could detect signals nearly 50 m away from the source. In another attempt, Arvind et al. (27) played back vocalizations of a rare bird in full volume on the speaker and determined the detection space to be around 700 m. Therefore, determining the detection space in PAM studies in the megadiverse and heterogeneous landscapes, such as India, is critical because it can improve the study design and optimize device deployment, resulting in robust inferences about biodiversity.

Much of India and other parts of South and Southeast Asia are extensively modified for agriculture. Despite intensification of agriculture, the landscapes comprise a heterogeneous mosaic of habitats and sustain a rich biodiversity (30). As part of our ongoing efforts to enhance biodiversity and encourage farmers to take up biodiversity-friendly agricultural practices, we initiated biodiversity assessments in rice paddy fields in Tamil Nadu state, South India. We found that agricultural fields are situated in a matrix of natural habitats such as forests and wetlands. Signals detected in the ARUs would thus be influenced by the combined effects of sound propagation, recorder sensitivity, directionality, as well as ambient masking noise beyond the physical attenuation of signals. Here, we present our findings from using experimental playback of known frequencies to determine the detection space of ARUs in rice paddy ecosystems. Specifically, we determine the change in detection probability across frequencies ranging between 1 – 8 kHz over a 50 – 300 m distance relative to the ARU and examine how the environmental variables affect them. In addition, we use a combination of four ARUs to determine how the direction of the source determines the probability of signal detection. Our study provides a crucial baseline and a pathway for rapidly determining the change in detection space across different habitats before initiating PAM to quantify and monitor biodiversity.

## 2 Materials and Methods

### 2.1 Study Area

We conducted this study in fallow agricultural fields outside the Kalakad Mundanthurai Tiger Reserve (KMTR, 8°25’ to 8°53’ N and 77°10’ to 77°35’ E) in Manimuthar Town, Ambasamudram Taluk, Tirunelveli District, Tamil Nadu, India. Sampling sites were chosen randomly in the Singampatti village (Table 1). We selected five agricultural fields in the post-harvest stage. Rice paddy is the predominant crop grown in this region, due to the availability of water from a network of canals during April/May – August/September and November – February, coinciding with the Southwest and Northeast monsoons respectively. This site is part of a larger study on biodiversity in agroecosystems, and therefore, the farmers were familiar with our work and facilitated the study. The paddy fields are typically between 0.5 – 1 acre in extent and are compartmentalized by a raised bund, which is used to hold water, as well as to walk to the paddy fields. The bunds, therefore, are typically devoid of shrubs, while occasionally, there may be trees planted. We chose sites at random relative to each other but ensured there was a 500 m straight line, without any major obstacles such as buildings, trees, or shrubs (Fig 1c).

**Fig 1:**
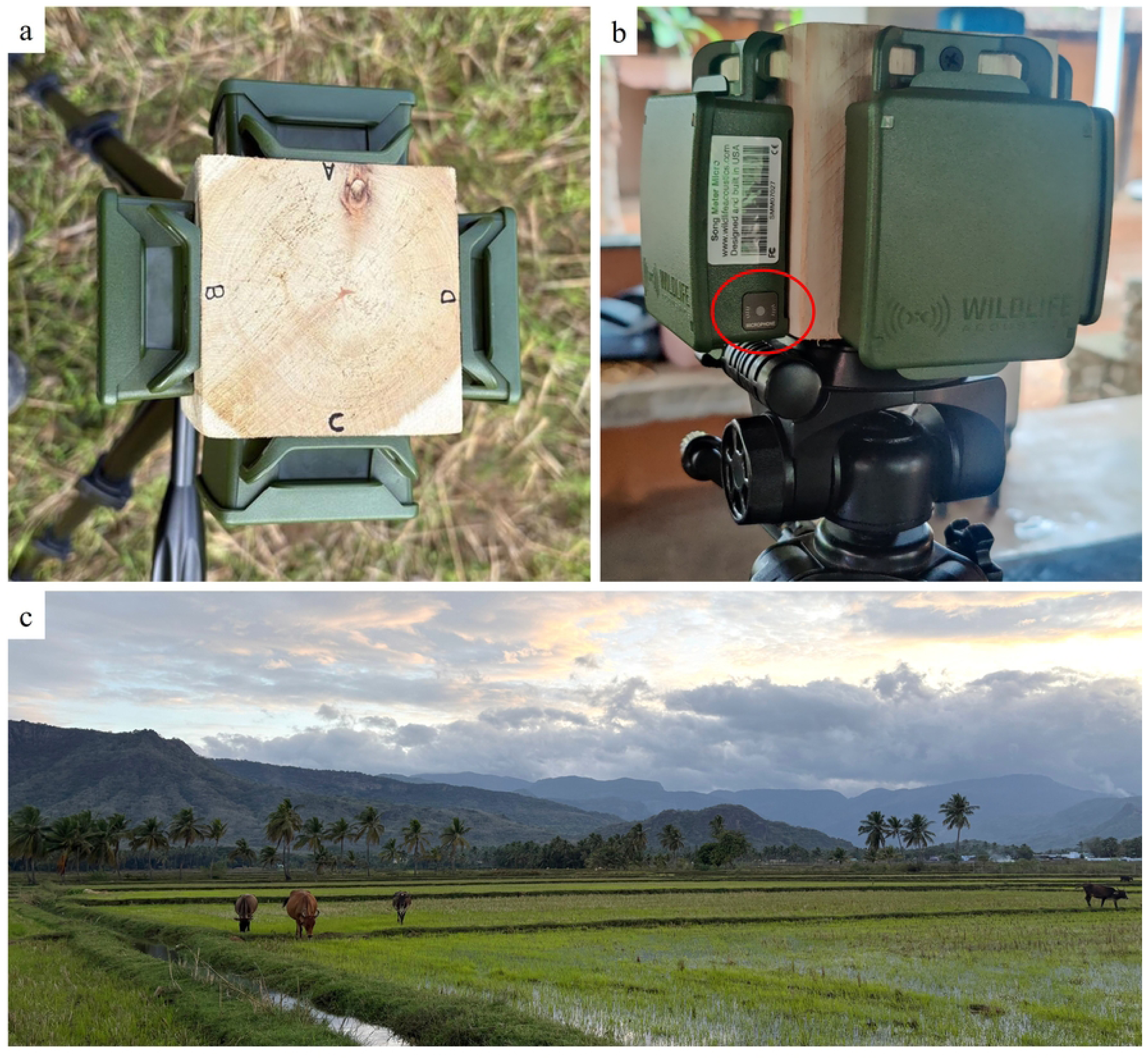
This study was conducted using four Song Meter Micro units. The four ARU were placed on a wooden block measuring 10 cm and mounted in four directions on a tripod (a). The position of the ARU was shuffled for each set of broadcast experiments. The ARU has one omnidirectional microphone, located on the right-hand side, indicated by the red circle (b). Broadcast experiments were conducted in fallow agriculture fields demarcated by the raised bund (c). The hills seen in the backdrop are part of the Kalakad Mundanthurai Tiger Reserve.

**Table 1:** Locations where signals were broadcast. All five locations are paddy fields.

| Name | Location.ID | Coordinates |
| --- | --- | --- |
| Therku Pappankulam | A | 8°39'16.2"N, 77°27'31.4"E |
| Zameen Singampatti | B | 8°39'33.5"N, 77°26'16.1"E |
| Singampatti | C | 8°39'17.9"N, 77°25'50.2"E |
| Manimuthar | D | 8°39'10.9"N, 77°25'08.1"E |
| Keezha Singampatti | E | 8°39'42.6"N, 77°26'24.5"E |

### 2.2 Study design

We relied on the playback of pure tone signals because it removes the variation in frequency and amplitude modulations in natural vocalizations of organisms (19). We generated five pure tone frequencies of 1, 2, 4, 6, and 8 kHz using Audacity^®^ 2.4.2. This range of frequencies covers the typical range of avian vocalizations at the study site, and the range allowed us to examine frequency-dependent detection space. Signals were broadcast using a portable public address megaphone (Ahuja Systems, PM 99, 25-Watt, battery-operated) at ∼1 m height, ensuring line-of-sight signal transmission to the receiver. Each playback was standardized to a sound pressure level (SPL) between 78 – 82 dB at 1 m from the speaker using a handheld sound level meter (Real Instruments HT 80A) at each distance and signal frequency. The speaker used was a directional horn type and was positioned facing the ARU, and the desired SPL was maintained by making slight adjustments to the input volume.

We used four Song Meter Micro units (Wildlife Acoustics Inc.), which feature a single integrated microphone located on the lower-right corner of the side. By mounting these in a fixed square configuration (Fig. 1a), we were able to assess how this hardware asymmetry, combined with device orientation, influenced signal reception. The recorder was set to record continuously with a sampling rate set to 44 kHz and a gain of 12 dB. We deployed four recorders simultaneously at ∼1 m height using a tripod with the following configuration: 0° (recorder facing the speaker), 180° (recorder facing in the opposite direction from the speaker), 90°, and 270° relative to the speaker to assess the directional response of the broadcast signals. Every day, the recorder orientation was rotated clockwise to randomize recorder angular response, if any. We recorded signals using this ARU setup at six linear distances from the speaker at each site: 50, 100, 150, 200, 250, and 300 m. We measured the distance using a measuring tape up to 50 m and subsequent distances using a laser rangefinder (Hawke LRF 400). We determined a 300 m cut-off based on initial attempts to find a straight, unobstructed paddy field and manual trials where three observers failed to detect the broadcast signal, and the visualization of recordings failed to show clear signals in the spectrograms.

### 2.3 Data Collection

We collected data between October 11-18, 2024, in two sessions: mornings (7:00 AM to 10:00 AM) and evenings (4:00 PM to 7:00 PM). We recorded the ambient environmental conditions at the start of each trial: Temperature (°C), relative humidity (%), and wind speed (km/hr) using a Kestrel Pocket Weather Meter 4500 (Nielsen-Kellerman^®^). Each trial consisted of 30 broadcast events (five frequencies × six distances). To organize the playback sequence and facilitate later identification of signals on spectrograms, we followed a consistent procedure for each frequency at a given distance. We played a brief 10 kHz cue (10 seconds) to mark the onset, then the target frequency tone was broadcast for 30 seconds, with a pause of 30 seconds to record ambient noise. After sequentially completing all five frequencies at the first distance (50 m), we moved to the next distance, repeating this process until a distance of 300 m. Across eight trials, we generated 240 broadcast events. As each event was to be recorded by all four ARUs, we expected a total of 960 recordings, but one ARU failed for a part of the study, and we only obtained 900 usable recordings. Broadcast trials were not conducted on the evenings of 11, 13, 14, and 17 October, the morning of 12, 14, 17, and 18 October due to equipment failure, rain, or both.

### 2.4 Acoustic Data Processing

We analyzed the audio recordings using Raven Pro V 1.6.5. Recordings were visualized as spectrograms using a Hann Window with a DFT size of 512 samples. Five independent observers, including the three who conducted the pre-experimental trials, reviewed all recordings using standard laptops and headphones while visually examining the spectrograms. Each recording was reviewed independently; a signal was marked as ’detected’ only if it was both audible to the observer and visible on the spectrogram. While signal-to-noise ratio (SNR) and in-band power are standard physical metrics for sound attenuation, they often fail to capture biological detectability in complex agroecosystems. In these landscapes, transient ambient noise from wind, insects, or machinery can fragment signals without necessarily rendering them undetectable to a trained observer or a classification algorithm. By focusing on an operative sense of detection, we directly model the real-world constraints of PAM workflows, where the primary goal is often species occupancy rather than quantifying physical attenuation. To incorporate the inherent uncertainty of this process, we recorded the number of positive detections (k) out of five for each playback. This count was modeled as a binomial response (k successes in 5 trials), which directly incorporates observer agreement and uncertainty.

### 2.5 Statistical Analysis

We modelled detection probability using a generalized linear mixed-effects model (GLMM) with binomial error distribution and logit link, using the ‘lme4’ package in R v4.4.2 (R Core Team 2024). For each playback event, the response variable was the number of observers detecting the signal out of five independent observer assessments, modelled as a binomial response. We included angle (relative to the broadcast source), time of the day, and frequency as factorial fixed effects. Distance to the source, ambient temperature, relative humidity, and wind speed were scaled and included as continuous fixed effects (scaled with mean = 0, SD = 1 to allow for better model convergence and interpretation); Location ID was included as a random factor to account for variation across experimental sites. To account for residual overdispersion, we included an observation-level random effect (ObsID, representing each unique recording) in the final model. The final model also included interaction terms for frequency × distance, frequency × temperature, frequency × humidity, and frequency × wind speed.

Sum contrasts were applied to all categorical predictors so that each estimate (reported on the log-odds scale) represents a deviation from the factor’s grand mean, rather than a specific reference level. Collinearity among predictors was checked using the Generalised Variance Inflation Factor (GVIF) via the car package, and all values were ≤ 2.1 . Model diagnostics were checked with the DHARMa package, and simulated residuals showed no significant deviation from uniformity (Kolmogorov–Smirnov test, p = 0.21) without overdispersion (dispersion test, p = 0.71) or zero-inflation (p = 0.93). Marginal probabilities were predicted and visualized using the ggeffects package. Marginal detection probabilities for recorder orientation were estimated using estimated marginal means derived from the fitted GLMM. Predictions were averaged over the levels of all other covariates and expressed on the response scale. Differences between ARU orientations were assessed using pairwise contrasts with Tukey adjustment for multiple comparisons.

## 3 Results

### 3.1 Factors affecting detection probability

Baseline detection probability varied significantly across the tested frequencies. Compared to the overall mean, the 1 kHz frequency had significantly lower baseline detection odds, whereas higher frequencies had positive coefficients (Table 2). Specifically, the 6 kHz signal had the highest increase in odds of detection, indicating it was detected most readily (Table 2). As expected, distance to the sound source exerted a strong negative effect on detection (p < 0.001, Table 2). However, this rate of decline was strongly frequency dependent. We observed a significantly sharper decrease for 1 kHz, and a positive interaction for 4 kHz, indicating the greatest retention of detectability over a distance gradient (Table 2). Overall, marginal probabilities indicate that higher frequencies maintained a higher effective detection probability than lower frequencies as distance to source increased (Fig 2). Among the environmental covariates, increased humidity significantly enhanced the likelihood of detection (Table 2). In contrast, the influence of ambient temperature, wind speed, and time of day (morning vs. evening) on the marginal probabilities were not statistically significant. However, a significant interaction between frequency of signal and temperature was observed. Increasing temperature significantly reduced the detection probability of the 1 kHz signal but significantly increased that of 2 kHz. Finally, baseline detectability was highly variable between sites used as a random effect (Variance = 0.95, SD = 1.0) as well as between observations (Variance = 15.0, SD = 3.8).

**Fig 2:**
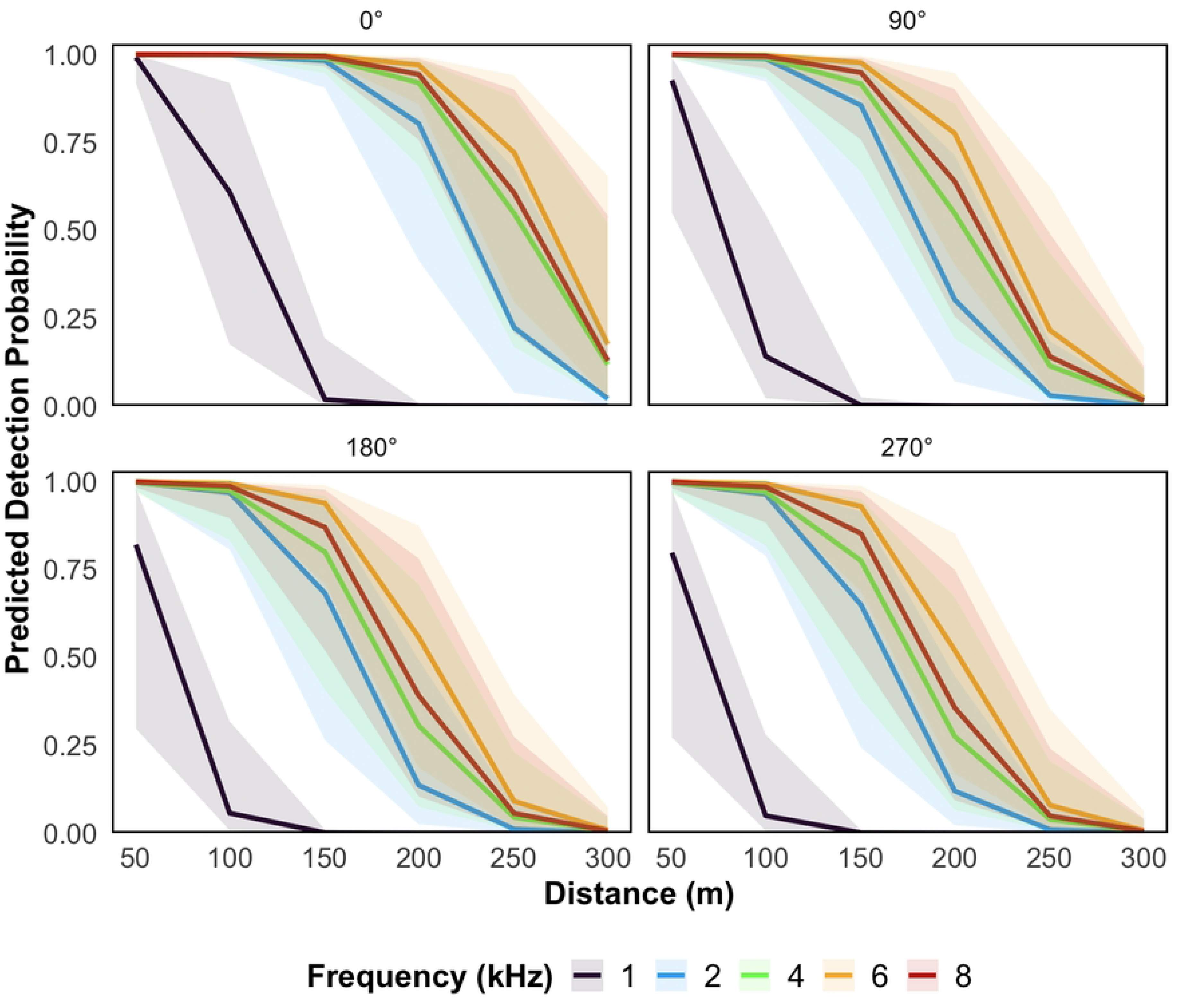
Marginal probabilities of detection across distances (50-300 m) for each broadcast frequency at all four angles decrease strongly with increasing distance. 4 kHz and 6 kHz showed the highest detection resilience at 0°, with 50% chances of detection even at ∼250-300 m. In contrast, 1 kHz showed the steepest decline in detectability.

**Table 2:** Fixed-effects summary from the binomial Generalized Linear Mixed-effects Model (GLMM) predicting the probability of acoustic signal detection. Predictors include categorical factors: frequency, recorder orientation, time of day, and scaled continuous covariates: distance, relative humidity (RH), temperature, and wind speed. Two-way interactions between signal frequency and all continuous covariates were modeled *apiori*. Estimates are reported on the log-odds scale utilizing sum contrasts for categorical variables. Statistically significant predictors are highlighted in bold (* p < 0.05, ** p < 0.01, *** p < 0.001).

| Variable | Estimate<br>(log odds) | Std. Error | z value | p value |
| --- | --- | --- | --- | --- |
| Intercept | -0.717 | 0.528 | -1.359 | 0.174 |
| **1 kHz** | **-7.906** | **0.938** | **-8.425** | **< 0.001\*\*\*** |
| **2 kHz** | **0.949** | **0.399** | **2.375** | **0.018 \*** |
| **4 kHz** | **1.772** | **0.393** | **4.503** | **< 0.001\*\*\*** |
| **6 kHz** | **2.967** | **0.407** | **7.282** | **< 0.001\*\*\*** |
| **Distance** | **-4.823** | **0.366** | **-13.190** | **< 0.001\*\*\*** |
| **RH** | **0.978** | **0.394** | **2.482** | **0.013 \*** |
| Temperature | 0.462 | 0.424 | 1.090 | 0.276 |
| WindSpeed | -0.145 | 0.264 | -0.550 | 0.582 |
| **0°** | **2.227** | **0.321** | **6.933** | **< 0.001\*\*\*** |
| 90° | -0.017 | 0.293 | -0.057 | 0.955 |
| **180°** | **-1.031** | **0.308** | **-3.344** | **0.001 \*\*** |
| Evening | 0.257 | 0.344 | 0.746 | 0.456 |
| **1 kHz × Distance** | **-2.654** | **0.846** | **-3.138** | **0.002 \*\*** |
| 2 kHz × Distance | 0.322 | 0.435 | 0.741 | 0.458 |
| **4 kHz × Distance** | **1.022** | **0.428** | **2.390** | **0.017 \*** |
| 6 kHz × Distance | 0.524 | 0.469 | 1.119 | 0.263 |
| 1 kHz × RH | -0.524 | 0.778 | -0.673 | 0.501 |
| 2 kHz × RH | 1.008 | 0.520 | 1.939 | 0.053 |
| 4 kHz × RH | 0.477 | 0.504 | 0.947 | 0.344 |
| 6 kHz × RH | -0.981 | 0.537 | -1.828 | 0.068 |
| **1 kHz × Temperature** | **-2.394** | **1.184** | **-2.023** | **0.043 \*** |
| **2 kHz × Temperature** | **1.426** | **0.501** | **2.847** | **0.004 \*\*** |
| 4 kHz × Temperature | 0.431 | 0.482 | 0.894 | 0.372 |
| 6 kHz × Temperature | 0.024 | 0.485 | 0.050 | 0.961 |
| 1 kHz × Wind Speed | -0.469 | 0.545 | -0.860 | 0.390 |
| 2 kHz × Wind Speed | -0.277 | 0.520 | -0.533 | 0.594 |
| 4 kHz × Wind Speed | 0.444 | 0.423 | 1.048 | 0.294 |
| 6 kHz × Wind Speed | 0.055 | 0.439 | 0.125 | 0.901 |

### 3.2 Effect of Recorder Orientation on Detection Probability

Recorder orientation relative to the sound source significantly influenced the odds of detection (Table 2). Marginal probabilities derived from the model (after holding all other covariates at their mean values) indicated that the ARU facing the sound source (0°) had an 81% probability of detection. In contrast, the detection probability for the ARU facing away from the source (180°) dropped to 14%. The lateral orientations (90° and 270°) exhibited intermediate marginal probabilities of 31% and 12%, respectively (Table 3). Post-hoc pairwise contrasts confirmed that the ARU facing the source (0°) had significantly higher odds of detection than all other orientations (Tukey adjusted p < 0.001; Table 4, Fig 3)

**Fig 3:**
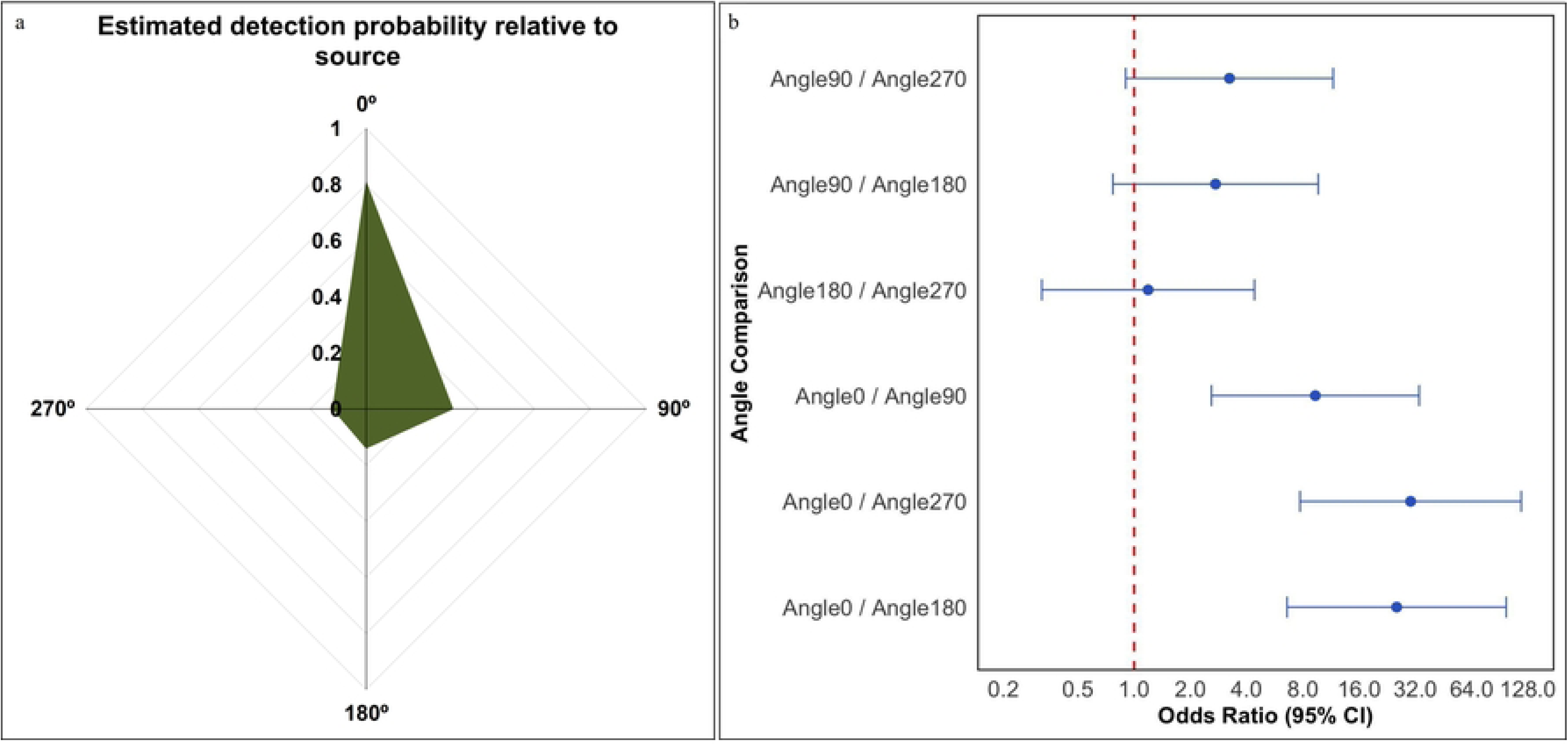
The placement of the recorder relative to the signal source influences detection drastically. If the recorder is placed facing the source, the detection probability is nearly 80% and drops if oriented away from the source (a). The odds also change when placed laterally (b). Each dot represents odds ratios (log10 scale) with 95% confidence intervals. The dashed red line at OR = 1 marks the threshold of no difference between angle pairs. Intervals that do not cross this line indicate statistically significant differences in detection odds between orientations (α = 0.05). The 0° (front-facing) recorder consistently outperformed other orientations.

**Table 3:** Modelled marginal probabilities of detection for each recorder orientation relative to the speaker.

| Angle | Modelled Marginal Probability | Std. Error | Lower 95% CI | Upper 95% CI |
| --- | --- | --- | --- | --- |
| 0° | 0.819 | 0.090 | 0.578 | 0.937 |
| 90° | 0.324 | 0.131 | 0.130 | 0.608 |
| 180° | 0.148 | 0.078 | 0.049 | 0.369 |
| 270° | 0.131 | 0.070 | 0.043 | 0.336 |

**Table 4:** Pairwise comparisons of detection probability across recorder orientations. An odds ratio greater than 1 indicates a higher detection probability for the first angle in the contrast compared to the second.

| Angle Comparison | Odds Ratio | Std. Error | z value | p value |
| --- | --- | --- | --- | --- |
| **Angle0 / Angle90** | **9.425** | **4.665** | **4.532** | **< 0.001\*\*\*** |
| **Angle0 / Angle180** | **25.992** | **13.561** | **6.244** | **< 0.001\*\*\*** |
| **Angle0 / Angle270** | **30.144** | **15.868** | **6.470** | **< 0.001\*\*\*** |
| Angle90 / Angle180 | 2.758 | 1.342 | 2.084 | 0.158 |
| Angle90 / Angle270 | 3.198 | 1.575 | 2.361 | 0.085 |
| Angle180 / Angle270 | 1.160 | 0.583 | 0.295 | 0.991 |

## 4 Discussion

Passive Acoustic Monitoring (PAM) using Autonomous Recording Units (ARUs) has emerged as a scalable solution for biodiversity assessment in recent years (31). Our study demonstrates that the ‘detection space’ or ‘active space’ of an ARU is highly frequency dependent. The 50% detection threshold for low-frequency signals (1 kHz) is approximately 100 m, whereas mid-range frequencies (∼6 kHz) remain reliably detectable up to 250 m. While signals are known to decay with increasing distance, environmental covariates further modulate these signals. Increased relative humidity significantly enhanced detection probability across the spectrum, while ambient temperature disproportionately influenced lower frequencies. Furthermore, we observed high site-level variability in baseline detectability between agricultural fields with visually similar habitat features (Table 5). In addition, the physical orientation of the ARU relative to the source exerted a dominant influence on performance, with only front-facing (0°) recorders achieving a high baseline detection of 81%. Taken together, these results demonstrate that without an empirical measure of detection space, PAM is prone to significant inferential bias. Failing to define such a detection space for any recorder leads to spatial over- or undersampling, which can erroneously skew species richness estimates. For instance, if used to gather evidence of restoration success, an ARU placed within a treatment plot may inadvertently record vocalizations from adjacent, untreated habitats such as a neighboring wetland or forest remnant. Without a calibrated detection space of ARUs, a researcher might attribute high acoustic activity to the success of a local intervention when the recorder ‘eavesdrops’ in the surrounding landscape or low success if the ARU fails to cover the intended study site. Such discrepancies restrict the comparability of monitoring efforts and risk misleading practitioners about the true effectiveness of habitat management for biodiversity conservation.

**Table 5:** Values are random intercepts (log-odds) for each Location ID from the binomial GLMM; positive values indicate higher baseline detection probability than the overall mean.

| Location ID | Baseline detectability<br>(log odds) |
| --- | --- |
| A | 0.071 |
| B | 1.084 |
| C | -0.462 |
| D | 0.284 |
| E | -1.238 |

### 4.1. Signal transmission and the environment

It is well known that acoustic signals in the field are shaped by a complex interplay of physical and environmental properties (32). The effectiveness of detecting acoustic signals depends entirely on accounting for signal masking and attenuation driven by vegetation structure, seasonality, and meteorological conditions (19,23,33). We provide the first systematic evidence that detection probability in tropical agroecosystems is not a linear function of distance, but a frequency-dependent detection space shaped by ambient masking. We observed a significantly steeper decline in detection probability for 1 kHz signals than for mid-range frequencies (4–6 kHz), which retained high detectability (> 50% probability of detection) up to 250 m. While lower frequencies are theoretically expected to carry further, our finding that 1 kHz signals diminished more rapidly at shorter distances is consistent with some of the earliest playback experiments (34) as well as recent ones (35), which show low frequency signals to decay faster, closer to the ground. This low detectability is likely driven by the ground effect and the specific soundscape of fallow paddy fields that acts as an acoustically soft surface (36). Therefore, the interference between direct and reflected waves from these surfaces can selectively damp certain frequencies depending on height and distance (37). In these open, human-modified landscapes, low-frequency ambient noise from wind, moving water in irrigation canals, distant anthropogenic activity, and atmospheric absorption also create a high masking floor affecting the propagation of all sound frequencies. As a result, low-frequency signals are the most affected, and we find an acoustic window between 4–6 kHz, where signals travel most efficiently with the least environmental interference.

Acoustic signals are also known to be affected by other environmental conditions such as relative humidity, wind speed, and temperature. Apart from masking, acoustic signals can also be absorbed, resulting in a loss of signal quality over distance. Snell-Rood (38) found the signal absorption to be non-linear and in warm tropical environments, attenuation peaks at low relative humidity (20–30%) and decreases with increasing relative humidity. Our findings are consistent with this pattern, where the higher levels of humidity positively affected the odds of detecting signals. In addition, we observed a complex interaction between temperature and low-frequency signal detection. Specifically, increasing temperatures significantly reduced the detectability of 1 kHz signals but had a positive effect on 2 kHz signals. This divergent response likely points to the microclimate at the air-ground interface in open fallow paddy fields. The negative impact on 1 kHz signals suggests that these longer wavelengths are particularly sensitive to this refractive loss and the convective turbulence associated with rising heat. In contrast, the positive response of 2 kHz signals may indicate that this frequency likely benefited from stabilized atmospheric conditions as the day progressed. These results underscore that the detection space of an ARU is not constant but dynamically responds to the environment. The high detectability we observed in the 4–6 kHz range is congruent with recent hardware evaluation by Mennill (35), who also found the Song Meter Micro to have peak sensitivity to middle frequencies with a reduced amplitude response at lower and upper ends of the frequency range between 1 – 20 kHz tested. We determined a frequency-dependent detection threshold (∼100 m for 1 kHz vs. ∼250 m for 6 kHz), which builds upon the work of Mennill (35), who found that while small ARUs are reliable within a 100 m radius for mid-frequency sounds, their effective detection radius is significantly constrained by both hardware sensitivity and environmental noise.

### 4.2. Directionality

Animal vocalizations can originate in any direction relative to the ARU. Thus, ARUs have an omnidirectional microphone and sample a notional sphere evenly around the microphone. However, we found little evidence of this assumption that omnidirectional microphones would detect signals evenly around them. We found a drastic decline in detection probability between the recorder facing the source (0°, 81%) and those facing away from the source (180°, 14%).

This pattern indicates the presence of an acoustic shadow, likely from the device itself and the mounting structure used. While Wildlife Acoustics (2023) specifies the Song Meter Micro microphone element as technically omnidirectional, our findings suggest that this characteristic is effectively nullified once the unit is mounted against a physical barrier. The distortion was most evident in the lateral asymmetry we observed (31% at 90° vs. 12% at 270°). The observed pattern is likely due to the position of the microphone on the ARU. The microphone is present on the bottom right-hand side of the housing, and when placed laterally to the sound source, the device body itself may cause an acoustic shadow. In our experimental configuration, the microphone of one lateral unit was effectively shielded by the corner of the wooden mounting block and the housing of the adjacent ARU (Fig 1a, b). What this means is that even when a recorder is oriented sideways to a sound source, its ability to detect signals is dictated by the location of the microphone on the device.

### 4.3. The necessity of local calibration for conservation monitoring

The direct implication of our findings for conservation practitioners would mean that a single ARU could effectively monitor different spatial scales simultaneously. A songbird vocalizing at ∼ 6 kHz might be sampled within a 250 m radius, while a dove vocalizing ∼1 – 2 kHz would only be detectable within a ∼50 – 100 m distance. Failing to account for these mismatched scales can lead to severe biases in species richness and community composition estimates (19,39). Beyond these frequency-dependent constraints, the physical deployment of the hardware further distorts signals. In most biodiversity monitoring programs, ARUs are strapped to thick tree trunks or wooden posts, which can cast a considerably larger acoustic shadow than our 10 cm experimental block. Therefore, any study using PAM should not assume an even 250 m radius sphere around the recorder, but account for the distortion by placing recorders to account for the acoustic data behind the ARU. We argue that omnidirectionality of the ARU should be treated as a theoretical maximum of microphone sensitivity rather than a field reality of the device. An easy solution to mitigate this issue would be to deploy ARUs in pairs, maintaining enough distance to avoid both underestimation and overestimation, or increase the sampling effort to account for deaf-spots, or orient the device towards high-priority habitat patches to ensure unbiased sampling. While such steps ensure unbiased sampling, they also double the capital expenditure and data processing efforts. Such a scenario highlights the ‘implementation gap’ between having affordable technology and having the resources to use it correctly (40).

Failing to account for these combined technical and biological limits of ARUs is evident in recent high-profile monitoring efforts. In India, PAM was used to determine the presence of the rare Jerdon’s Courser (*Rhinoptilus bitorquatus*). Although the researchers determined the detection space to be up to 800 m, it was based on call playbacks at maximum speaker output volume (27). While such tests define technical capability, they do not reflect biological reality. By standardizing our playbacks to a biologically realistic 80 dB (at 1 m), we demonstrate that a 50% detection threshold of 100–250 m is a more conservative and reliable estimate for survey design. Relying on maximum-volume calibration risks a substantial overestimation of the monitored area, potentially leading to false absences if recorders are spaced apart. Perhaps the subsequent detection of the bird after several decades underscores this point (BirdLife International 2025). Furthermore, PAM is increasingly proposed for evaluating ecological restoration success (28). However, without accounting for frequency-dependent decay and orientation bias, richness estimates in restoration plots may be confounded by signal spillovers from adjacent high-quality habitats. In mosaic landscapes, an uncalibrated ARU may ‘eavesdrop’ into a neighboring forest, recording high richness erroneously attributed to the local restoration intervention. This ‘eavesdropping’ is a significant inferential threat when using community-wide metrics like acoustic indices.

### 4.4. A Proposed Decision Support Framework for PAM

The rapid expansion of PAM has often outpaced the development of rigorous survey protocols, resulting in a situation where the collected data may not be informative for management (40). While workflows already exist (15,40), none include the estimation of detection space despite acknowledging that detection radii are not even across devices. To bridge this gap, we propose a ‘Decision Support Framework’ that integrates technical and environmental constraints from our empirical findings (Fig 4). This framework operates as a sequential, five-stage process:

**Fig 4.**
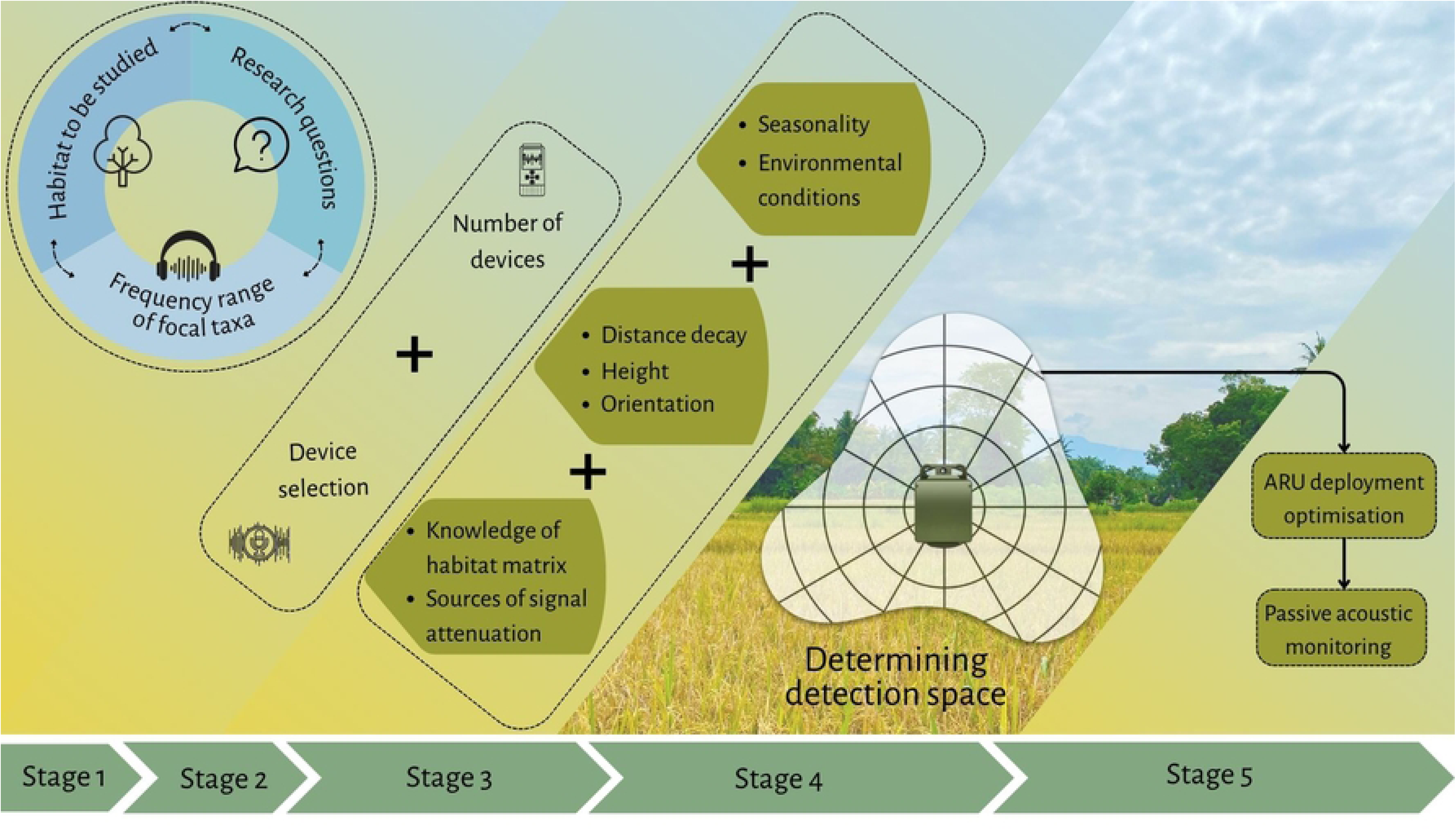
A five-stage decision-support framework for optimizing Autonomous Recording Unit (ARU) deployment in Passive Acoustic Monitoring (PAM). The sequential workflow moves from iterative project planning (Stage 1) and hardware logistics (Stage 2) to the integration of physical, environmental, and habitat-specific calibration parameters (Stage 3). Rather than assuming a uniform circular radius, these parameters are synthesized to determine the true, often asymmetrical, detection space of the recorder (Stage 4). By combining this empirically defined space with device availability, practitioners can optimize field deployment (Stage 5) to mitigate acoustic shadows and signal spillover.

Stage 1: Conceptualization and Focal Taxa: Researchers must first establish the habitat to be studied, the core research questions, and the focal taxa, which directly dictate the target frequency range. This forms an iterative planning loop, as the frequency range of the focal taxa itself limits the types of ecological questions that can be reliably answered using PAM.

Stage 2: Hardware Logistics: Based on the requirements established in Stage 1, practitioners finalize device selection and assess their logistical constraints (i.e., the total number of devices available).

Stage 3: Calibration Parameters: Researchers must evaluate the specific limiting factors of their study site. This involves integrating physical acoustics (distance decay, device height, and hardware orientation), atmospheric modulators (seasonality and environmental conditions), and the surrounding habitat matrix (which dictates sources of signal attenuation and potential signal spillover).

Stage 4: Determining Detection Space: Synthesizing the variables from Stage 3, researchers empirically or theoretically map the actual ‘active space’ of the recorder. As our results demonstrate, this space should not be assumed to be a notional circular grid, but rather an irregular envelope constrained by the environment and hardware acoustic shadows.

Stage 5: Deployment Optimization and Execution: The empirically determined detection space is combined with the logistical constraints of device availability (from Stage 2). This ensures that the final ARU deployment optimizes landscape coverage while actively mitigating hardware blind spots and eavesdropping, culminating in robust Passive Acoustic Monitoring.

To operationalize this framework, we provide a practical deployment matrix based on our GLMM marginal probabilities (Table 6). By selecting their target taxa’s peak frequency and their desired statistical confidence (e.g., high-confidence inventory vs. baseline occupancy), practitioners can extract the recommended inter-recorder spacing and mandatory hardware orientation. For example, ensuring 90% detection confidence for a 1 kHz target requires <50 m spacing between the recorder and the species, along with a strict 0° orientation toward the habitat. In contrast, a 6 kHz target permits a wider 150 m grid. As India scales its national PAM networks (15), adopting such a calibrated, framework-driven approach will be essential for turning acoustic recordings into robust, defensible conservation policy.

**Table 6.**
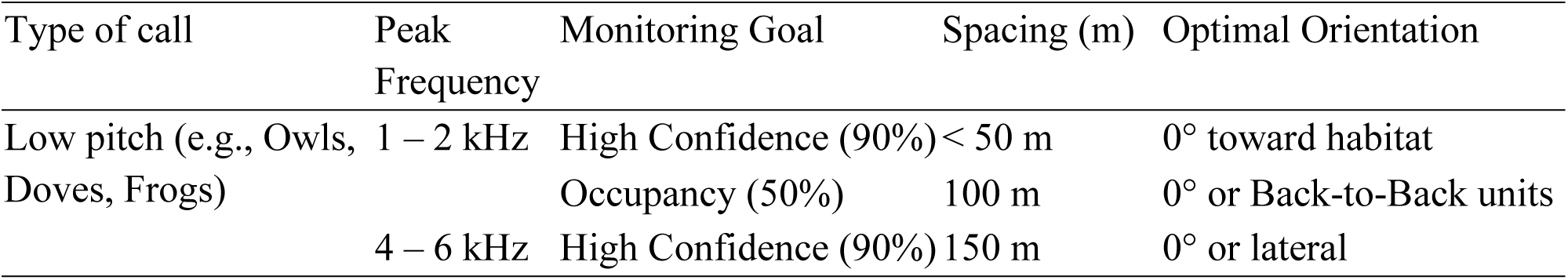
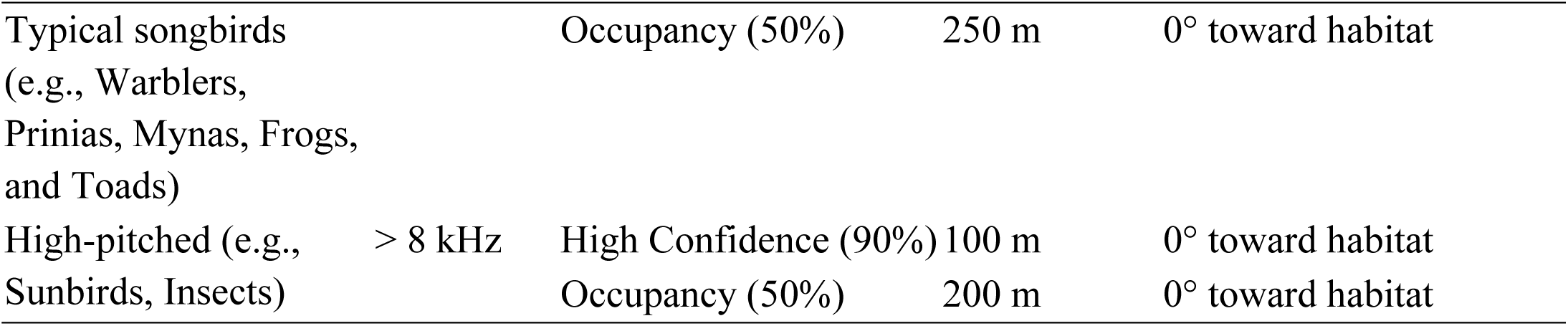
Suggested framework for deploying ARU in agroecosystems. 0° refers to the front of ARU facing the source.

### 4.5 Future Directions

While our study provides a foundational calibration for PAM in tropical agroecosystems, several factors warrant further investigation to refine these detection models. First, we relied on pure tone signals to allow for a controlled estimation of frequency-specific decay and recorder sensitivity. While this method has been used (35), using animal vocalizations instead can capture the effects of complex frequency and amplitude modulations, varying durations, and harmonic structures. Second, we did not explicitly quantify the radiation pattern of the broadcast speaker. While playbacks were standardized to a fixed SPL, some portion of the observed orientation effect may reflect source directionality in addition to the recorder’s angular response. Given that many vocalizing organisms exhibit high source directionality, future studies should aim to characterize the synergistic effects of both source and receiver orientation. Our results clearly indicate that the omnidirectionality of the ARU is a theoretical maximum rather than a field reality. Thus, accounting for the acoustic spread of both signalers and the sensitivity of sensors will be critical for high-precision density estimation. Third, we relied on detections in the recordings as a measure, rather than parameters such as signal-to-noise ratios. Although signal-to-noise ratio (SNR) and in-band power are commonly used to quantify acoustic signal attenuation, these metrics were not used here because they were not consistently well-defined under field conditions with the equipment available to us. Ambient noise varied substantially across trials, and at larger distances, signals were often intermittently visible or fragmented in spectrograms, making continuous amplitude-based measurements unreliable. Instead of modeling change in amplitude with distance for each frequency, we then focused on whether signals were detected, using multiple independent observer assessments, which allowed us to incorporate uncertainty directly, rather than discarding ambiguous cases or relying on noisy amplitude data. This approach of focusing on effective detectability considers SNR, device response, and ground effect that reflects the way PAM is used in biodiversity monitoring. For instance, in determining the richness of species, the key question is “was the species detected?” rather than “what was the received level?” Finally, the temporal scope of this study was limited to the post-harvest season. Tropical agroecosystems are highly dynamic; detection space likely fluctuates significantly across seasons due to changes in vegetation height (monsoon growth), ambient noise profiles (choruses and rain), and the microclimate. While wind speed was not a significant factor in our specific trials, the high variability of wind in open landscapes during different seasons may alter the masking and disproportionately affect low-frequency detection. Similarly, our findings on the positive influence of humidity on signal detection suggest that the effective monitoring radius expands and contracts across a 24-hour cycle. Future calibration efforts should prioritize these seasonal and diel fluctuations to ensure that long-term PAM datasets remain comparable and robust.

## 5 Conclusions

Our study provides the first systematic calibration of Passive Acoustic Monitoring (PAM) in Indian agroecosystems, revealing that the effective detection space of a recorder is a dynamic volume shaped by signal frequency, hardware orientation, and environmental conditions. We demonstrate that detection space is not a uniform sphere, but a distorted envelope constrained by an 86% data loss in the rear-facing hemisphere. This acoustic shadow effectively refutes the field-level assumption of omnidirectionality, underscoring the importance of hardware placement and mounting geometry as critical to data quality as the sensitivity of the microphone itself. Furthermore, we identify an acoustic window in the 4–6 kHz range where signals travel most efficiently, contrasting with a high masking floor and ground effect, which severely limits the detection of low-frequency taxa to within 100 m. Without the frequency-specific calibration curves provided here, researchers risk eavesdropping into adjacent high-quality habitats or undersampling, resulting in erroneous estimates, especially in restoration plots or agricultural interventions. We propose a Decision Support Framework for ARU deployment, based on our findings. By integrating focal taxa traits with technical and environmental constraints, researchers can move from notional monitoring to evidence-based survey designs that are robust and comparable across landscapes. As India and other megadiverse nations scale their national PAM networks, adopting these calibration standards will be essential for ensuring that acoustic data are translated into defensible conservation policy and effective land management.

## Author Contribution

PS: Data Curation, Validation, Formal Analysis, Software, Visualization (lead), Writing-Original Draft, Writing-Review and Editing

KK: Conceptualization, Investigation, Methodology, Writing-Review and Editing

KSS: Conceptualization, Supervision, Methodology, Funding Acquisition, Project Administration, Visualization, Writing-Review and Editing

## Acknowledgements

This study was part of the field immersion component of the ATREE-TDU MSc in Conservation Practice Course, and the following students assisted in data collection: Harshin Meera Ambavarapu, Hassan Shahnowaz Islam, Rutu Paresh Bhanushali, Niharika Gowda, and Priya Dharshini Venkatachalam. The staff members of the Agasthyamalai Community Conservation Reserve, Manimutharu, provided invaluable field support and assistance. The farmers of Manimutharu permitted us to conduct playback in their paddy fields. Dr Anand Krishnan and Arpit Omprakash provided valuable insights and fruitful discussions on earlier versions of the MS. Vidisha M K provided illustrations and reviewed the MS. This work was funded by the generous support of the Dhanam Foundation and the DST Inspire Faculty Fellowship to KSS.

